# A new multi-genomic approach for the study of biogeochemical cycles at global scale: the molecular reconstruction of the sulfur cycle

**DOI:** 10.1101/148775

**Authors:** Valerie De Anda, Icoquih Zapata-Peñasco, Bruno Contreras-Moreira, Augusto Cesar Poot-Hernandez, Luis E. Eguiarte, Valeria Souza

## Abstract

Despite the great advances in microbial ecology and the explosion of high throughput sequencing, our ability to understand and integrate the global biogeochemical cycles is still limited. Here we propose a novel approach to summarize the complexity of the Sulfur cycle based on the minimum ecosystem concept, the microbial mat model and the relative entropy of protein domains involved in S-metabolism. This methodology produces a single value, called the Sulfur Score (SS), which informs about the specific S-related molecular machinery. After curating an inventory of microorganisms, pathways and genes taking part in this cycle, we benchmark the performance of the SS on a collection of 2,107 non-redundant RefSeq genomes, 900 metagenomes from MG-RAST and 35 metagenomes analyzed for the first time. We find that the SS is able to correctly classify microorganisms known to be involved in the S-cycle, yielding an Area Under the ROC Curve of 0.985. Moreover, when sorting environments the top-scoring metagenomes were hydrothermal vents, microbial mats and deep-sea sediments, among others. This methodology can be generalized to the analysis of other biogeochemical cycles or processes. Provided that an inventory of relevant pathways and microorganisms can be compiled, entropy-based scores could be used to detect environmental patterns and informative samples in multi-genomic scale.

## INTRODUCTION

Despite their fundamental importance in sustaining life on Earth, understanding the fluxes of fundamental elements (C, H, O, N, S, and P) through the Earth’s surface has been challenging for several reasons (Li *et al.*, 2012; Newman and Banfield, 2002). First, the global biogeochemical cycles are enormously complex as they are an interconnected network of biological, chemical and geophysical processes that have been coevolving in the biosphere since the apparition of the first metabolic processes on Earth (~3.8 billion years ago) (Falkowski *et al.*, 2008). Since then, the evolutionary history of life on Earth has been shaped by a complex synergistic cooperation of multispecies a*ss*emblages that differ in terms of ecological and metabolic capabilities (Canfield 2005). Secondly, although these assemblages are often spatially and temporally separated, they effectively couple electron transfer (i.e., redox) reactions that transform elements and energy derived from several abiotic processes (Falkowski *et al.*, 2008). Thirdly, these abiotic processes involve the continuous supply and removal of elements from various Earth surface reservoirs, such as geothermal processes derived from mantle and crust, tectonics and rock weathering, and photochemical processes in the atmosphere including the constant flux of solar energy (Hedges, 1992).

As a result of these challenges, the fluxes of fundamental atoms through Earth have been studied and approached using different disciplines that address specific layers of complexity. For instance, geochemistry and atmospheric sciences have been focused on addressing the major abiotic process at global planetary scales, i.e., processes influencing the flux of elements to and from the various Earth surface reservoirs and atmosphere (Canfield et al., 2005, Falkowski *et al.*, 2008).

Moreover, microbial ecology has emphasized in understanding the links between microbial catalyzed activities and ecosystem and biogeochemical processes (Morales and Holben, 2011). Current approaches for establishing metabolic relationships *in situ* are based on targeting coding-sequences involved in specific biochemical pathways related to carbon, nitrogen, sulfur, and phosphorus cycling. In this way, DNA or RNA extracted directly from natural environments is sequenced and quantified with conventional PCR and Sanger sequencing or microarray analyses (Loy *et al.*, 2004; Khodadad and Foster, 2012) for example the GeoChip (Tu *et al.*, 2014); or other ‘omics’ techniques such as metagenomics (Quaiser *et al.*, 2011; Swingley *et al.*, 2012; Llorens-Marès *et al.*, 2015) or metatranscriptomics (Stewart *et al.*, 2011; Chen *et al.*, 2015).

However, despite the great advances in microbial ecology and high throughput sequencing, our ability to understand and integrate the global biogeochemical cycles is still limited. Here, we propose a new comprehensive approach to evaluate, compare, and facilitate the study of such cycles based on the metabolic machinery of the microorganisms responsible for driving element fluxes throughout the Earth. The approach is based mainly on three aspects: i) the minimum ecosystem concept: which considers the properties, forces (outside energy), flow pathways (energy and matter), interactions, and feedback loops or circuits for the flow of matter or energy (Odum,1993); ii) the microbial mats, which are nearly closed and self-sustaining ecosystems that comprise the major biogeochemical cycles, trophic levels and food webs in a vertical laminate pattern (Bolhuis *et al.*, 2014); and iii) the mathematical rationalization of Kullback-Leibler divergence, also known as relative entropy *H’* (Kullback and Leibler, 1951). Relative entropy has been widely applied in physics, communication theory and statistical inference, and is interpreted as a measure of disorder, information and uncertainty, respectively (Commenges, 2015). Here we use the communication theory concept of H’ to summarize the information conveyed by the protein domains (metabolic machinery) encoded by environmental DNA sequences. The application of this measure in biology was originally developed by Stormo and colleagues to identify binding sites that regulate gene transcription sites (Hertz and Stormo, 1999).

In this context, the compartmentalization of microbial mats provides clear, natural or arbitrary boundaries that evoke the concept of “minimum ecosystem”, which can be delimited in a natural or arbitrary sense (Odum, 1993). Therefore, specific parts of the cycle may be seen as parts of a whole. For instance, the redox level, reduced-oxidized compounds or even genes and enzymes implicated in certain routes can be used as ecosystem boundaries. These assemblies set up a unit that represents the minimum ecosystem with the minimum requirements to be functional, therefore this can be applied to measure the information derived from the complexity inside any biogeochemical cycle.

To test and evaluate the performance of our conceptualization, we focused on the biogeochemical Sulfur cycle (from now on S-cycle), due to its critical role in the biogeochemistry of the planet - *i.e.*, respiration of sedimentary organic matter, oxidation state of the atmosphere and oceans, and the composition of seawater (Halevy *et al.*, 2012). Despite the extensive biochemical knowledge of both oxidative and reductive microorganisms (Rabus *et al.*, 2013; Canfield, 2015; Dahl, 2017), there are no studies aiming to integrate all the microbiological and geochemical transformations and their corresponding metabolic pathways of the sulfur cycle. Our study proposes a general computational approach that can be easily modified and used to compare and measure other biogeochemical cycles. This procedure generates measurable scores to evaluate these cycles and their importance and ecological weight on a global scale.

## MATERIALS AND METHODS

The computational analysis addressing the different levels of complexity of the S-cycle was divided into four stages illustrated in Figure 1. The corresponding scripts and documentation are available for download from: https://github.com/eead-csic-compbio/metagenome_Pfam_score.

**Figure 1.**
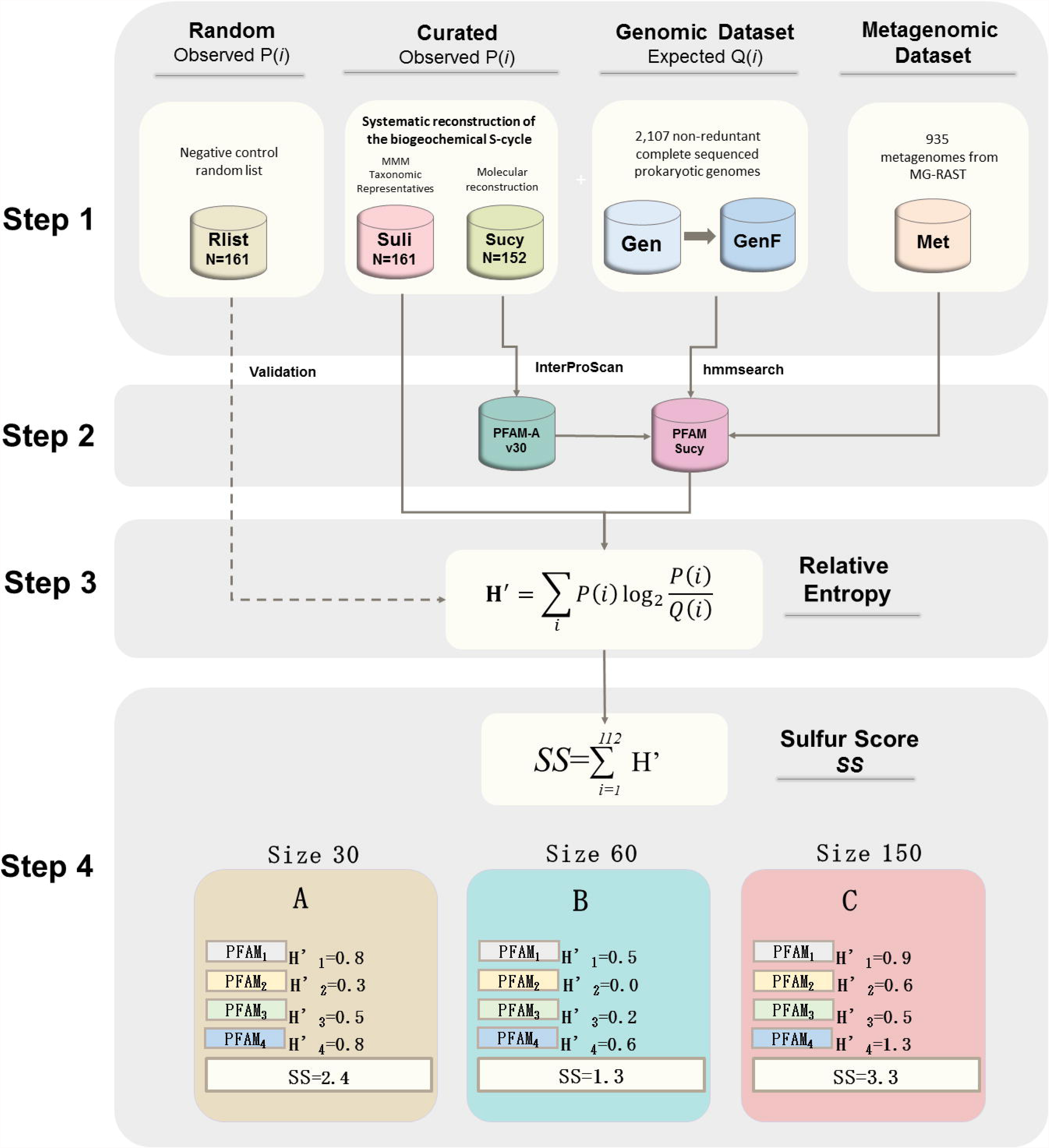
Computational pipeline for the analysis of the different levels of complexity of the sulfur cycle (S-cycle). The first step is to obtain the datasets. The biogeochemical processes of the S-cycle were compiled in two lists. First, the most important S-compounds involved in abiotic or abiotic reactions were curated to produce a database of metabolic pathways, redox reactions, enzymatic numbers and their corresponding coding genes (Sucy). Second, a list of S-based microorganisms (Suli) was also curated and compiled using the information of the Microbial Mat Model described in Figure 2. Then a total of 1,000 list of random genomes were used as negative control (Rlist). The information gathered was then used to evaluate the ‘omic’ datasets (Gen, GenF, Met). Stages 2 to 4 summarize the determination of the Sulfur Score (*SS*) with sets of peptide sequences of increasing mean size length (MSL, in this example A, B and C). *SS* is calculated by summation of the entropy of 112 scanned protein domains, each one with a certain entropy value (*H’*).

### STAGE 1: The biogeochemical complexity of S-cycle and ‘omic’-datasets

#### Taxonomic representatives of sulfur cycle: the microbial mat model

According to the minimum ecosystem concept, we consider microbial mats as models of a minimum ecological system (Microbial Mat Model). Based on the metabolic guilds found in microbial mats and other S-derived environments (i.e., hot springs, black smokers, sludge, etc.), we reviewed primary literature and the MetaCyc database (Caspi *et al.*, 2012) to select S-based microorganisms (at genus or species level) with experimental evidence of the physiology of degradation, reduction, oxidation, or disproportionation of organic/inorganic S-compounds. The list of S-based prokaryotes is found in Table S1. The non-redundant list of these S-based microorganisms with fully sequenced genomes (December 2016) was called the Sulfur list (Suli), which currently contains 161 genomes used as input of the pipeline.

#### Random taxonomic representatives (RList)

In order to build negative control sets of organisms that are not particularly enriched on metabolic preferences, 1,000 random lists of microorganisms were drawn using the genomic dataset explained below as reference, with the same number of microorganisms included in Suli.

#### Metabolic pathways and genes

We gathered and classified the metabolic pathways involved in the S-cycle from the primary literature and expert-curated databases KEGG (Kanehisa and Goto, 2000) and MetaCyc (Caspi *et al.*, 2012). This molecular information was integrated into a single database named Sulfur cycle (Sucy), which currently contains 152 genes and 48 enzyme classification numbers annotated in the Enzyme classification (http://enzyme.expasy.org) (Table S2). The 152 FASTA sequences of the peptides encoded by these genes were downloaded from UniProt (Magrane and Consortium, 2011) and used as the input of the pipeline.

#### Omic datasets

In order to test, compare and evaluate the importance of the S-cycle in ‘omic’ datasets, we generated the following databases:

i. *Genomic dataset* (Gen): Due to the redundancy of complete genomes deposited in RefSeq (https://www.ncbi.nlm.nih.gov/genome/browse/reference; 4,158 genomes at the time of the analysis, December 21, 2016), we decided to reduce the set of genomic data by using a web-based tool, that uses “genomic similarity scores” (Moreno-Hagelsieb *et al.*, 2013). Selecting values of genomic similarity of 0.95 and a DNA signature of 0.01, we obtained a total of 2,107 non-redundant genomes in FASTA format.
ii. *Metagenomic dataset* (Met): We selected metagenomes from the MG-RAST server version 3.6 that met the following conditions: i) publicly available; ii) metadata associated; and iii) environmental samples (isolated from defined environments o features like rivers, soil, biofilms), discarding microbiome host-associated metagenomes (*i.e.*, to human, cow, chicken). In addition, we also included 35 unpublished metagenomes derived from sediment, water and microbial mats from Cuatro Ciénegas (Coahuila, Mexico), which were also submitted to the MG-RAST pipeline. Using the above-mentioned conditions, a total of 935 metagenomes were downloaded in FASTA format (http://api.metagenomics.anl.gov/api.html, coding regions within the reads, December, 2016). Then, we measured the Mean Size Length (MSL) of the peptide coding regions of the Met FASTA files (Figure S1A). Taking into account that the 152 sulfur proteins (Sucy) have lengths ranging from 49 to 1,020 amino acid residues (aa), their detection in metagenomic samples will likely be affected by MSL (Figure S1B). For example, the identification of long proteins harboring several domains (i.e., catalytic, cofactor binding etc.) might be impaired in metagenomes with short MSL.
iii. *Fragmented genomic dataset* (GenF). We used the genomic dataset as a reference for benchmarking the detection limits of the protein families. The peptide FASTA-format files of the 2,107 non-redundant genomes were *in silico* sheared into seven categories of increasing fragment length, taking into account the variation in read sizes of the metagenomic dataset (Figure S1A).

### STAGE 2: Domain composition of the sulfur proteins

We used Interproscan 5.21-60.0 (Jones *et al.*, 2014) to annotate the protein domains encoded in the 152 Sucy genes, according to the Pfam-A database v30 (Finn *et al.*, 2008). In total, 112 Pfam domains where identified and subsequently scanned against the ‘omic’ datasets with HMMER 3.0 (Finn *et al.*, 2011).

### STAGE 3: Relative entropy and its use to detect informative sulfur-related protein domains

In order to obtain an estimate of how protein families are represented in S-based microorganisms, we used a derivative of the Kullback-Leibler divergence (Kullback and Leibler, 1951) — also known as relative entropy *H’(i)* — to measure the difference between probabilities *P* and *Q* (see Eq. 1 below). In this context, *P(i)* represents the frequency of protein domain *i* in Suli genomes (observed frequency), while *Q(i)* represents the frequency in the complete genomic dataset (expected frequency). *H’*, in bits, captures the extent to which a domain informs specifically about sulfur metabolism. *H*’ values that are close to 1 correspond to the most informative Pfam domains (enriched among S-based genomes), whereas low *H’* values (close to zero) describe non-informative ones. Negative values correspond to proteins observed less than expected.

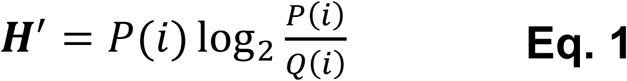

As a control, *H’* was recalculated both in Gen and GenF datasets replacing Suli with 1,000 equally sized lists of random-sampled genomes (Rlist). Using these procedures, we evaluated the variation of relative entropy of each Pfam domain in order to i) short-list those that could be used as markers in metagenomic dataset (regardless of their MSL) and ii) to generate a score which could be used to compare the importance of S-metabolism in ‘omic’-samples (either in Gen or Met).

#### Clustering of Pfam domains according to their entropy

The following clustering algorithms were tested: K-Means, Affinity propagation, Mean-shift Spectral, Ward hierarchical, Agglomerative, DBSCAN and Birch. These methods are part of the scikit-learn Machine Learning Python module (http://scikit-learn.org/stable/modules/clustering.html).

### STAGE 4: The Sulfur Score (*SS*): origin, interpretation, properties and benchmark

We propose to evaluate the importance of biogeochemical S-cycle in ‘omic’-datasets using a single metric that we call “Sulfur Score” (*SS*) (Eq. 2). By this approach, sulfur informative protein domains would contribute to higher *SS*, whereas non-informative ones would decrease it. This is an extension of procedures originally developed for the alignment of DNA and protein motifs, in which individual positions are independent and additive, and can be simply summed up to obtain the total weight or information content (Hertz and Stormo, 1999). Instead of aligning sequences, in our context we compare a presence/absence string of Pfam domains, from which a total weight (*SS*) is computed.

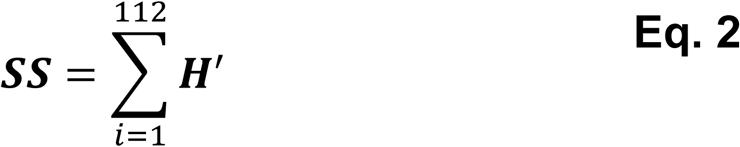

If we compare total *SS* across several genomes or environments, those in which the majority of metabolic pathways of S-cycle are represented will thus have a high *SS*; in contrast, low *SS* values should be expected if proteins involved in the S-cycle are not particularly enriched.

#### Calibration

We took into account the MSL of each metagenome to compute the *SS*. Briefly; we gathered the entropy values (*H’*) of the 112 Pfams in Gen and GenF (Figure S2A). *H’* estimates vary among the different GenF categories, with the major differences observed for fragments of 30 and 60 aa (Figure S2B) and estimates converging with increasing MSL. Therefore, the *SS* algorithm considers the MSL of the ‘omic’-sample and chooses the appropriate baseline *H’* values pre-computed on the GenF dataset. The GenF fragment size ranges (30, 60, 100, 150, 200, 250 and 300) match the observed MSL in real metagenomic sets (see Figure S1).

#### Properties and performance of SS

Because the outcome of the *SS* depends on i) the list of S-related Pfam domains and ii) the curated list of S-genomes, we measured its reproducibility with several approaches. First, scores computed in 2014 (Pfam v27, 1528 non-redundant genomes, 156 species in Suli) were compared to the current results (Pfam v30, 2017 non-redundant genomes, 161 curated species in Suli). Second, we compared the outcomes of the *SS* using a random sampling experiment. Briefly, we computed *SS* 1000 times both in the Gen and the Met datasets sorted in terms of their GenF size category. In each test, ≈50% of the 112 S-related Pfam domains were randomly selected to compute *SS*. Finally, we benchmarked the predictive capacity of the *SS* in order to accurately classify the genomes of S-related organisms (Suli, n=161, positive instances) in contrast with a larger set of non-redundant genomes (Gen ^-^Suli, n=1.946, negative instances). True Positive Rates (TPR), False Positive Rates (FPR) and the resulting of Receiver Operating Characteristic (ROC) plots were produced with the scikit-learn module (http://scikit-learn.org/stable/modules/model_evaluation.html), and finally the Area Under the ROC Curve (AUC) was computed.

## RESULTS AND DISCUSSION

### Defining the biogeochemical S-cycle

What parameters define a biogeochemical cycle, and what are its limits and scope? Which elements should be considered necessary for each cycle? The study of element fluxes between rocks, atmosphere, oceans and biological activity can be extremely complex in terms of space and time, ranging from single living cells to entire ecosystems, and can be completed in microseconds or instead developed over geological time scales (Hedges, 1992; Falkowski *et al.*, 2008; Zhao *et al.*, 2014; Olson *et al.*, 2016; Widder *et al.*, 2016). Currently, microbialecology techniques are insufficient to capture the integral complexity of the biogeochemical cycles by selecting a few marker genes to evaluate the importance of any given element in the environment. Here, we argue that a comprehensive description of the relationship between complex biotic an abiotic process is crucial to describe and understand the importance of global biogeochemical cycles and provide a method to do so by taking advantage of ‘omic’-era data.

We propose a new approach to analyze, compare, evaluate and quantify the importance of biogeochemical cycles in ‘omic’ datasets summarized in Figure 1, focusing, as a case-study, on the S-cycle. The first step consists on the systematic acquisition of the molecular and ecological information required to describe the cycle of interest. This information can be considered an inventory (Odum, 1993).

With the manual curation effort, we obtained both: a list of microorganisms (Suli), and a list of genes encoding enzymes (Sucy) related to the S-cycle. The Suli list includes the microorganisms involved in the global S-cycle using the microbial mats as ecological model. Microbial mats are compartmentalized organizations that were the first ecosystems to appear on Earth and have evolved over more than three billion years into the complex ecosystems that we know today (Herman EK and Kump LR, 2005). Functionally, microbial mats are self-sufficient structures that support most of the major biogeochemical cycles within a vertical dimension of only a few millimeters in a multilayer space (Pinckney and Paerl, 1997) (Figure 2A). These assemblies represent ecosystems with the minimum requirements to be functional and therefore can be used to explore the complexity of biogeochemical cycles. In contrast with the compact nature of microbial mats, the distribution of the metabolic S-guilds is widely dispersed at planetary scale, being found in rivers and estuaries, lagoons, oceans, sediments and deep hydrothermal vents (Halevy *et al.*, 2012).

**Figure 2.**
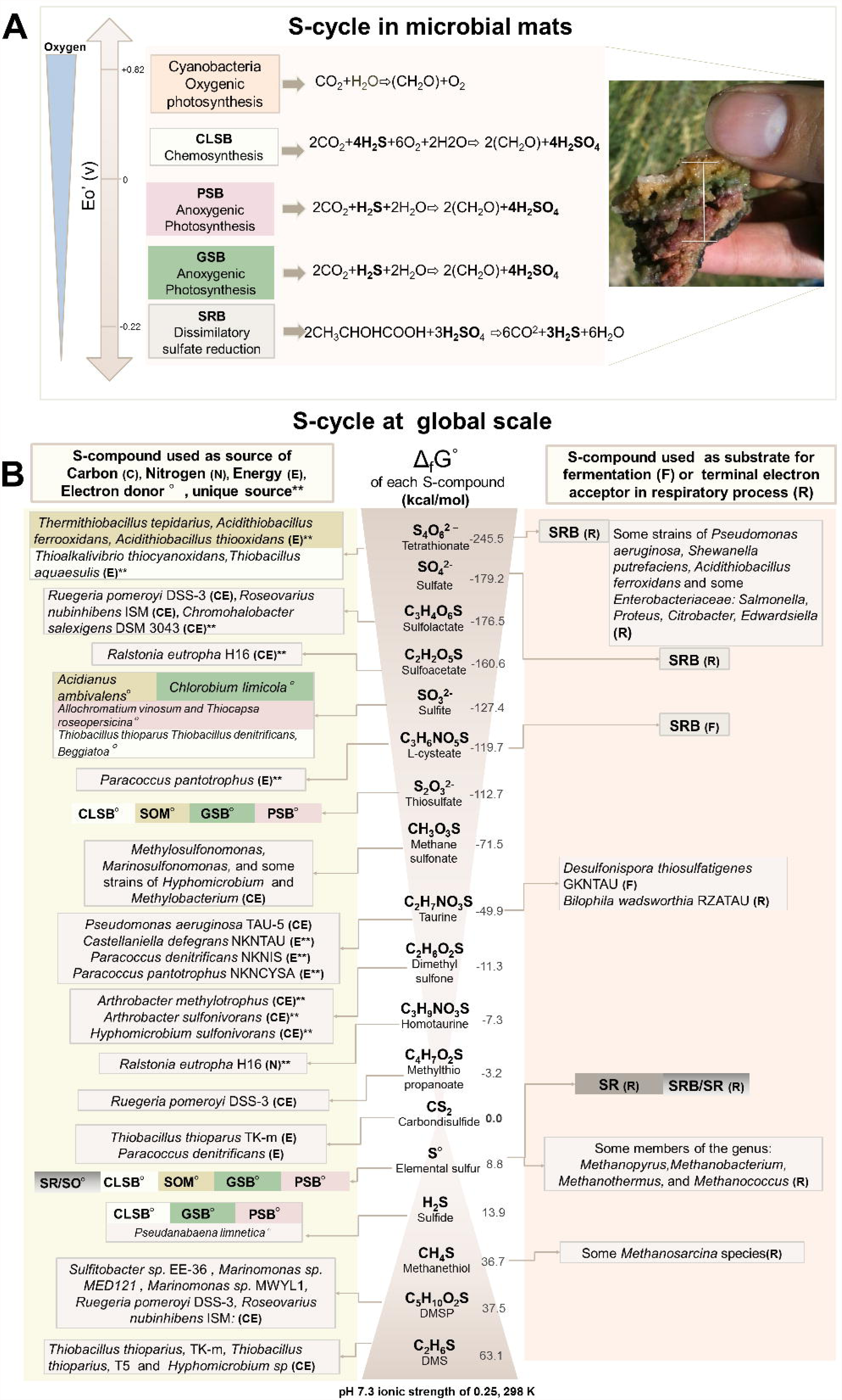
Sulfur cycle in small and planetary scale. A) Microbial Mat Model: Simplified scheme of the relevant reactions carried out in microbial mats according to the redox potential. The redox couples (at pH 7) are approximate; the exact values depend upon how the reactions are coupled. 1) oxygenic photosynthesis by Cyanobacteria 2) chemosynthesis by chemolithotrophic <u>c</u>olor-<u>l</u>ess <u>s</u>ulfur <u>b</u>acteria (CLSB), anoxygenic photosynthesis by 3) <u>p</u>urple <u>s</u>ulfur <u>b</u>acteria (PSB) and 4) green <u>s</u>ulfur <u>b</u>acteria (GSB). sulfate reduction performed by 5) <u>s</u>ulfate <u>r</u>educing <u>b</u>acteria (SRB). B) Sulfur cycle at planetary scale. Most important organic and inorganic S-compounds derived from biogeochemical processes arranged according to the Standard Gibbs free energy of formation described in Caspi *et al.*, (2012). The left column indicates whether specific microorganisms are able to use those S-compounds, as a source of Carbon (C), Nitrogen (N), Energy (E) or Electron donors (°). Double asterisks indicate if the S-compound is used as sole source, of C, N, or E. The corresponding electron acceptors in redox-coupled reactions using the S-compound as electron donor are not show. The right column indicates whether the S-compounds are used as fermentative substrate (F) or terminal electron acceptor in respiratory processes (R). Colored boxes summarized the metabolic guilds involved in the metabolism of S-compounds, in oxidation (*i.e.*, CLSB, SOM, PSB, and GSB) or reduction (SR, SRB) process. Some redox coupling reactions carried out by the latter metabolic guilds are showed in panel A. The complete list of S-based microorganisms (Suli) is found in Table S1. Figure based on annotations from MetaCyc (Caspi *et al.*, 2012).

The distribution of S-related bacterial taxa can be analyzed in terms of redox potential and Gibbs Energy of Free Formation of S-compounds, resembling the compact layout of the metabolic guilds in the microbial mat (Figure 2B). Therefore, Suli includes three main groups of microorganisms belonging to the S-metabolic guilds in microbial mats i) chemolithotrophic color less sulfur bacteria (CLSB), ii) anaerobic phototrophs: purple sulfur bacteria (PSB), green sulfur bacteria, (GSB), and iii) sulfate reducing bacteria (SRB) as well as deep-branch sulfur hyperthermophile microorganisms found in extreme conditions (hot springs, black smokers, etc.), such as elemental sulfur reducing (SRM) and oxidizer (SO) microorganisms.

The other manually curated list of the inventory, Sucy, contains the metabolic pathways, genes and enzyme activity numbers involved in the S-fluxes (Table S2). To the best of our knowledge, this is the first attempt to integrate the biotic and abiotic processes involved in the mobilization of inorganic/organic S-compounds through microbial-catalyzed reactions at global scale; we gathered and classified all the metabolic pathways involved in the S-cycle according to three key aspects described in Figure 2B, i) S-compound: either organic or inorganic, derived from abiotic or biotic processes, ii) standard Gibbs free energy of formation (GFEF), and iii) metabolic role of the S-compound. The metabolic pathways involved in these S-compounds were systematically divided into 28 pathways (Table S3). We included the pathways involved in a) the oxidation/reduction of inorganic S-compounds, used as source of energy, electron donor or acceptor, b) the degradation of organic S-compounds such as aliphatic sulfonates sulfur amino acids, and organosulfonates, c) methanogenesis from methylated thiols such dimethyl sulfide DMS, metylthio-propanoate and methanethiol, which are generated in nature by different biogeochemical processes (Caspi *et al.*, 2012), and d) the biosynthesis of sulfolipids (SQDG), because it has been observed that some bacteria living in S-rich and P-lacking environments, are able to synthetized sulfolipids instead of phospholipids in the membrane as an adaptation of the selective pressures of the environment (Alcaraz *et al.*, 2008).

Once we integrated the metabolic inventory (genes, enzymatic numbers and major metabolic complexes), we linked all the enzymatic steps and S-compounds into a comprehensive representation of the S-cycle in a single cell (Figure 3), with the following features i) the comprehensive interconnection of pathways in terms of energy flow, ii) the direction of the reactions of the important biogeochemical S-compounds, iii) the interplay of the redox gradient (organic/inorganic) of the intermediate compounds that act as key axes of organic and inorganic reactions (i.e., sulfite), and iv) the molecular reconstruction of S-cycle at different levels (genes, abiotic sulfur-derived compounds and enzymatic steps).

**Figure 3.**
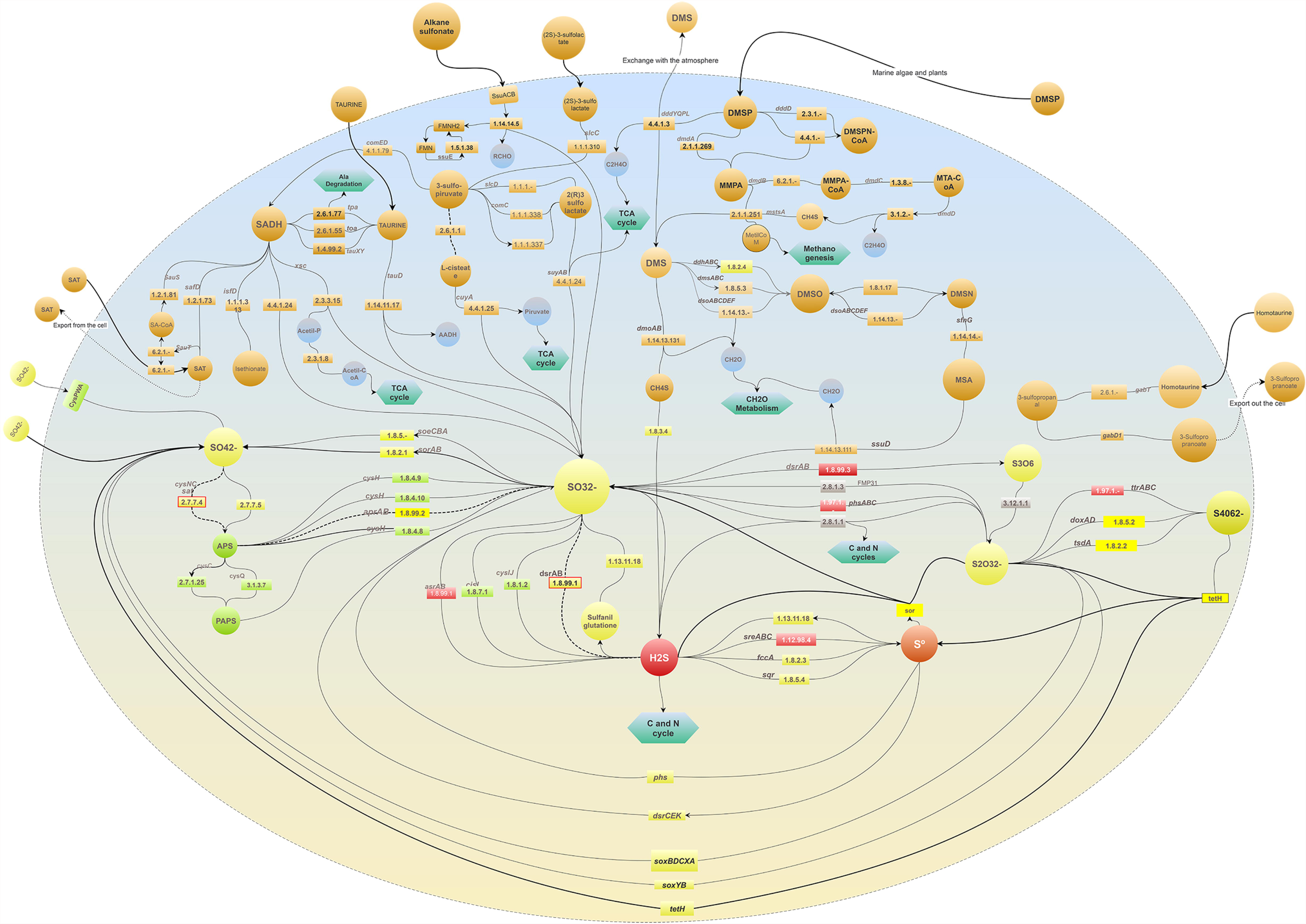
Comprehensive representation of the global biogeochemical S-cycle assembled from many metabolic pathways found in a variety of organisms combined in a single cell. To the best of our knowledge, all the molecular pathways involved in the metabolism of sulfur compounds, described in Figure 2B, are included. The enzymatic steps are depicted as rectangles followed by arrows indicating the direction of the reaction. Green hexagons represent metabolic links to other metabolisms. Bold dashed arrows indicate bidirectional reactions. Inorganic S-compounds have been arranged according to their reduction potential, from the most oxidized (yellow) to the most reduced (red) compounds. Grey rectangles indicate enzymes acting in disproportionation processes in which a reactant is both oxidized and reduced in the same chemical reaction, forming two separate compounds. Input biogeochemical S-compounds are shown outside and connected with bold arrows. Dashed arrows indicate S-compounds excreted out of the cell. The upper half of the modelled cell depicts the processes involved in the use of organic S-compounds (orange circles) found in natural environments and used as source of carbon, sulfur and/or energy in several aerobic/anaerobic strains described in Figure 2.

In order to benchmark the entropy-based approach described below, we used available data in both genome and metagenome databases. We also included 35 unpublished metagenomes derived from microbial mats, water and sediment from an ultra-oligotrophic shallow lake in Cuatro Cienegas, Coahuila (CCC), Mexico. Altogether, these 935 metagenomes set up the Met dataset. The Gen dataset was sheared with different fragment sizes taking into account the Mean Size Length (MSL) of Met (Figure S1), producing dataset GenF, which was used for estimating the detection limits of sulfur protein families. We describe the computation approach step-by-step below.

### Relative entropy as a way to identify protein domains that inform about the S-cycle

After the first step of collecting the datasets, the second step involved the annotation of the coding sequences of the 152 genes in Sucy, yielding a total of 112 Pfam domains (Pfam Sucy in Figure 1). The third step consisted in measuring the relative entropy (*H’)* of each Pfam using Equation 1. The presence of each Pfam domain in Suli (genome list) and in the genomic datasets (Gen and GenF) was used as observed and expected frequencies respectively. The obtained *H’* values are shown in Figure S2.

As negative control, we tested to what extent those *H’* values depend on the particular genomes curated in Suli. To do so, we replaced Suli by 1,000 lists of random-sampled genomes and used them to compute the observed frequencies. As expected, there was a clear difference between the *H’* computed using random genomes (Figure S3A), and those obtained with the manually curated list of S-based microorganisms (Figure S3B). In particular, entropy values derived from the random test were found to be approximately symmetric and consistently low among the GenF size categories, yielding values of -0.09, and 0.1 as 5% and 95% percentiles, respectively (Table S4).

We then evaluated the behavior of the *H’* values in both Gen and GenF to test whether informative Pfam domains can be used as molecular markers of S-cycle in metagenomic sequences of variable MSL. In order to be considered as a marker gene, each Pfam domain had to fulfill three requirements 1) produce consistently high mean *H’* values in Gen and GenF, 2) display low standard deviation (std), and 3) obtain *H’* values clearly separated from the random distribution. We tested several clustering methods, summarized in Figure S4; among them, the Birch and Ward methods grouped together the informative protein domains with low std (Figure S5). However, Ward clustering included a few protein domains relevant in the S-cycle, which were otherwise discarded by the Birch method. Therefore, we selected the Ward method, which produced three clusters of protein domains described in Figure 4.

**Figure 4.**
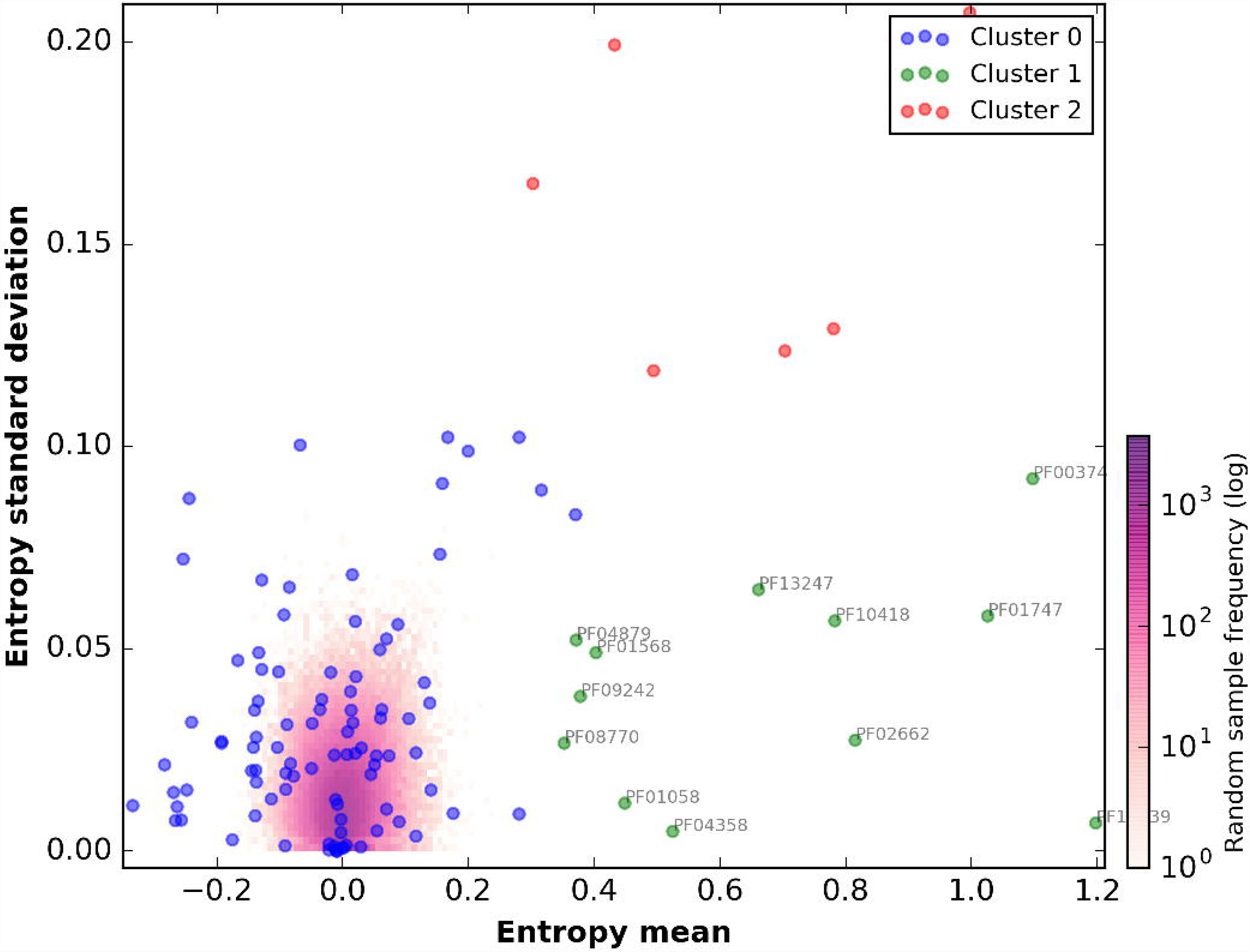
Clustering of the Pfam relative entropies obtained in Gen and GenF produced with the Ward method. Log frequency of the entropy values computed in the random test is colored in purple (see scale bar). Cluster 0 (blue) groups protein domains with low relative entropy that overlap with the random distribution. Cluster 1 (green) includes the Pfam domains that fulfill the requirements to be used as molecular markers (high H’ and low standard deviation, std). Red dots (cluster 2) correspond to Pfam domains with high H’ and std.

Cluster 0 includes 94 domains with entropies in the range [-0.4, 0.4], thus overlapping those obtained in the negative control explained earlier. Cluster 1 identifies 12 Pfam domains listed in Table 1, with high entropy and low std, and can therefore be proposed as molecular markers. Among the proposed molecular marker protein domains are APS-Reductase (PF12139: *H’*=1.2), ATP-sulfurilase (PF01747: *H’*=1.03) and DsrC (PF04358: *H’*=0.52), key protein families in metabolic pathways involved in both sulfur oxidation/reduction processes. Finally, cluster 2 groups 6 domains (described in Table S5) with high entropy values and high std, such as PUA-like domain (PF14306: *H’=1*). We presume that the protein domains listed in Table S5 are key players in S-metabolism and their presence in almost all complete-sequenced S-associated microorganisms suggests their important roles.

**Table 1.**
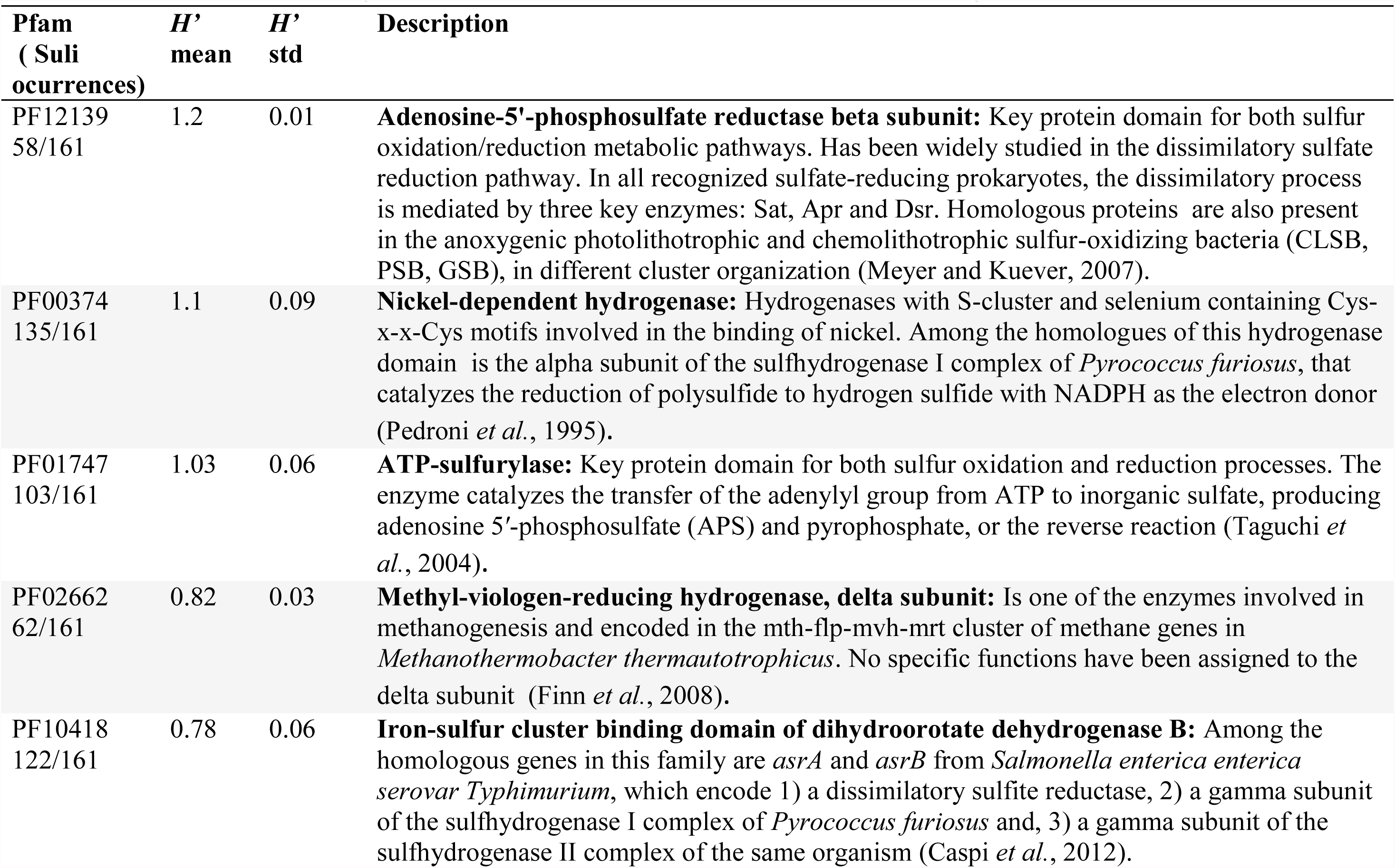
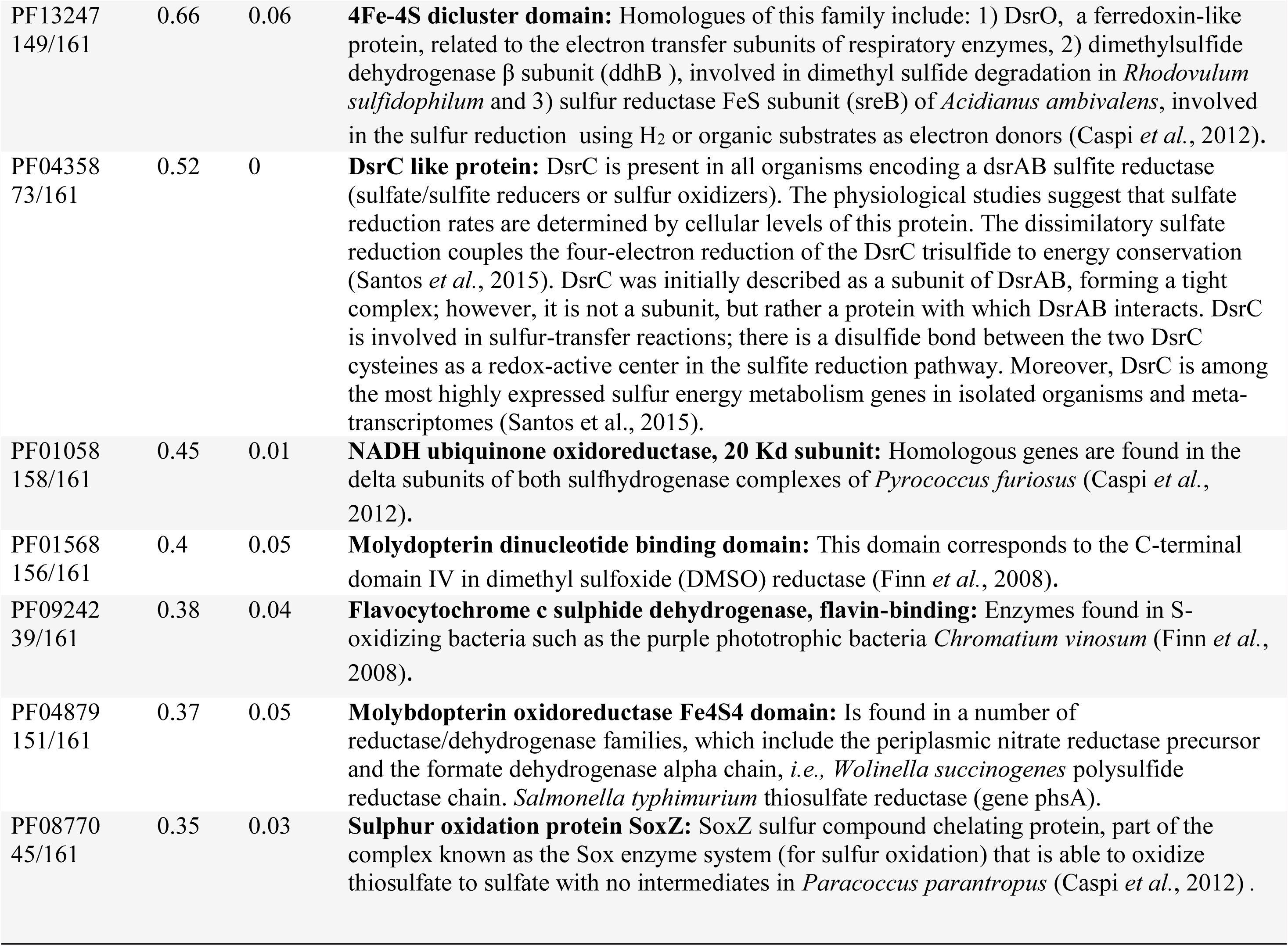
Informative Pfam domains with high H’ and high std. Novel proposed molecular marker domains in metagenomic datasets of variable mean size length (MSL)

| Pfam<br>( Suli<br>occurrences) | $H'$<br>mean | $H'$<br>std | Description |
| --- | --- | --- | --- |
| PF12139<br>58/161 | 1.2 | 0.01 | **Adenosine-5'-phosphosulfate reductase beta subunit:** Key protein domain for both sulfur oxidation/reduction metabolic pathways. Has been widely studied in the dissimilatory sulfate reduction pathway. In all recognized sulfate-reducing prokaryotes, the dissimilatory process is mediated by three key enzymes: Sat, Apr and Dsr. Homologous proteins are also present in the anoxygenic photolithotrophic and chemolithotrophic sulfur-oxidizing bacteria (CLSB, PSB, GSB), in different cluster organization (Meyer and Kuever, 2007). |
| PF00374<br>135/161 | 1.1 | 0.09 | **Nickel-dependent hydrogenase:** Hydrogenases with S-cluster and selenium containing Cys-x-x-Cys motifs involved in the binding of nickel. Among the homologues of this hydrogenase domain is the alpha subunit of the sulfhydrogenase I complex of *Pyrococcus furiosus*, that catalyzes the reduction of polysulfide to hydrogen sulfide with NADPH as the electron donor (Pedroni *et al.*, 1995). |
| PF01747<br>103/161 | 1.03 | 0.06 | **ATP-sulfurylase:** Key protein domain for both sulfur oxidation and reduction processes. The enzyme catalyzes the transfer of the adenylyl group from ATP to inorganic sulfate, producing adenosine 5'-phosphosulfate (APS) and pyrophosphate, or the reverse reaction (Taguchi *et al.*, 2004). |
| PF02662<br>62/161 | 0.82 | 0.03 | **Methyl-viologen-reducing hydrogenase, delta subunit:** Is one of the enzymes involved in methanogenesis and encoded in the mth-flp-mvh-mrt cluster of methane genes in *Methanothermobacter thermautotrophicus*. No specific functions have been assigned to the delta subunit (Finn *et al.*, 2008). |
| PF10418<br>122/161 | 0.78 | 0.06 | **Iron-sulfur cluster binding domain of dihydroorotate dehydrogenase B:** Among the homologous genes in this family are *asrA* and *asrB* from *Salmonella enterica enterica* serovar *Typhimurium*, which encode 1) a dissimilatory sulfite reductase, 2) a gamma subunit of the sulfhydrogenase I complex of *Pyrococcus furiosus* and, 3) a gamma subunit of the sulfhydrogenase II complex of the same organism (Caspi *et al.*, 2012). |
| PF13247<br>149/161 | 0.66 | 0.06 | **4Fe-4S dicluster domain:** Homologues of this family include: 1) DsrO, a ferredoxin-like protein, related to the electron transfer subunits of respiratory enzymes, 2) dimethylsulfide dehydrogenase $\beta$ subunit (ddhB), involved in dimethyl sulfide degradation in *Rhodovulum sulfidophilum* and 3) sulfur reductase FeS subunit (sreB) of *Acidianus ambivalens*, involved in the sulfur reduction using $H_2$ or organic substrates as electron donors (Caspi *et al.*, 2012). |
| PF04358<br>73/161 | 0.52 | 0 | **DsrC like protein:** DsrC is present in all organisms encoding a dsrAB sulfite reductase (sulfate/sulfite reducers or sulfur oxidizers). The physiological studies suggest that sulfate reduction rates are determined by cellular levels of this protein. The dissimilatory sulfate reduction couples the four-electron reduction of the DsrC trisulfide to energy conservation (Santos *et al.*, 2015). DsrC was initially described as a subunit of DsrAB, forming a tight complex; however, it is not a subunit, but rather a protein with which DsrAB interacts. DsrC is involved in sulfur-transfer reactions; there is a disulfide bond between the two DsrC cysteines as a redox-active center in the sulfite reduction pathway. Moreover, DsrC is among the most highly expressed sulfur energy metabolism genes in isolated organisms and meta-transcriptomes (Santos *et al.*, 2015). |
| PF01058<br>158/161 | 0.45 | 0.01 | **NADH ubiquinone oxidoreductase, 20 Kd subunit:** Homologous genes are found in the delta subunits of both sulphydrogenase complexes of *Pyrococcus furiosus* (Caspi *et al.*, 2012). |
| PF01568<br>156/161 | 0.4 | 0.05 | **Molybdopterin dinucleotide binding domain:** This domain corresponds to the C-terminal domain IV in dimethyl sulfoxide (DMSO) reductase (Finn *et al.*, 2008). |
| PF09242<br>39/161 | 0.38 | 0.04 | **Flavocytochrome c sulphide dehydrogenase, flavin-binding:** Enzymes found in S-oxidizing bacteria such as the purple phototrophic bacteria *Chromatium vinosum* (Finn *et al.*, 2008). |
| PF04879<br>151/161 | 0.37 | 0.05 | **Molybdopterin oxidoreductase Fe4S4 domain:** Is found in a number of reductase/dehydrogenase families, which include the periplasmic nitrate reductase precursor and the formate dehydrogenase alpha chain, *i.e.*, *Wolinella succinogenes* polysulfide reductase chain. *Salmonella typhimurium* thiosulfate reductase (gene phsA). |
| PF08770<br>45/161 | 0.35 | 0.03 | **Sulphur oxidation protein SoxZ:** SoxZ sulfur compound chelating protein, part of the complex known as the Sox enzyme system (for sulfur oxidation) that is able to oxidize thiosulfate to sulfate with no intermediates in *Paracoccus parantropus* (Caspi *et al.*, 2012). |

Despite their different properties, all the 112 Pfam domains mentioned in those clusters were subsequently used in the next sections to detect peptides related to the S-cycle in ‘*omic*’ datasets.

### Identification of S-based genomes using the Sulfur Score

To test whether Pfam entropies can be combined to capture the S-related metabolism, we calculated the Sulfur Score (*SS*) with Equation 2 for all non-redundant genomes in dataset Gen. The obtained *SS* and the corresponding taxonomy according in NCBI for each genome can be found in Table S6. Then we classified all the genomes according to their metabolic capabilities in three subsets: Suli) containing manually 161 curated genomes; Sur) Sulfur unconsidered or related microorganisms not included in Suli with *SS* > 4 (in total 192), which were curated afterwards, and NS) including 1,754 Non-Sulfur species, comprehends the subset Gen – (Suli + Sur). Boxplots summarizing the scores for these subsets are plotted in Figure 5A.

**Figure 5.**
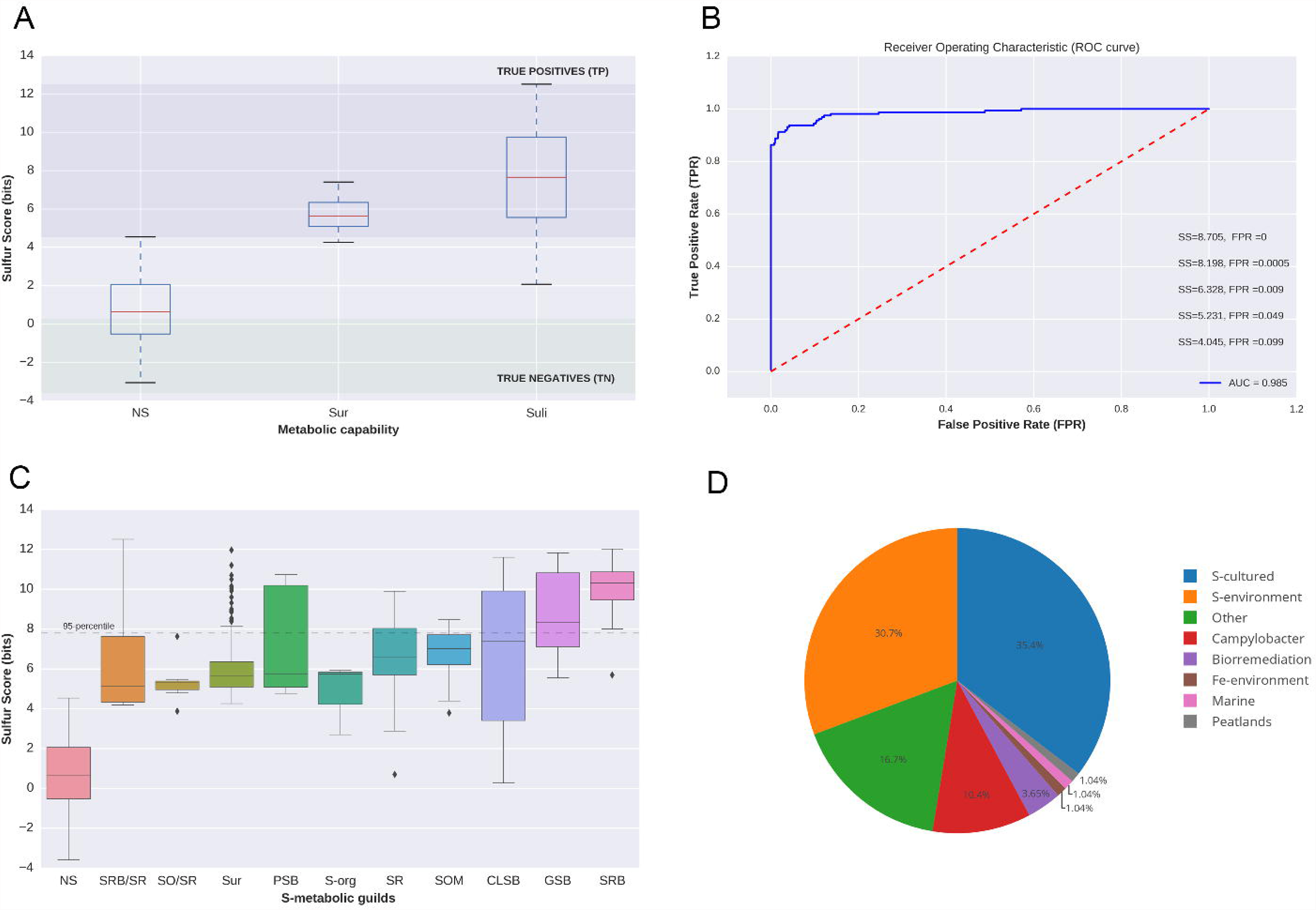
Distribution of Sulfur Score (*SS*) in genomes of dataset Gen. A) Subsets of non-redundant genomes: i) genomes annotated in Suli (n=161); ii) Sur, genomes not listed in Suli with *SS* > 4 and candidates to be S-related microorganisms (n=192); iii) rest of the genomes in Gen (NS, n=1,754). According to the curated species, True Positives can be defined as genomes with SS > max (SSNS) distribution, whereas True Negatives are those with SS < min(SSS_uli_). B) ROC curve with Area Under the Curve (AUC) indicated together with thresholds for some False Positive Rates (FPR). C) Distribution of SS for different S-metabolic guilds according to the microbial mat model (Figure 2A) and also the genomes in Sur. D) Assignment of the 192 genomes in Sur to ecological categories based on literature reports.

In order to measure the predictive value of *SS*, we computed a Receiver Operator Characteristic (ROC) curve by calling positive instances those annotated in Suli and negative the rest of the genomes, while iterating along increasing values of *SS*. The results are shown in Figure 5B, with an estimated Area Under the Curve (AUC) of 0.985, and the cut-off values of *SS* for several False Positive Rates (FPR). With this training Gen dataset, *SS*=8.705 is required to rule out all false positives. However, *SS*=5.231 is sufficient to achieve a FPR < 0.05. Figure 5C breaks down the species in Suli according to the metabolic guilds of the microbial mat model observing the clear difference between the distribution of *SS* in NS and Sulfur-related genomes (Suli and Sur). Finally, the *SS* values of candidate genomes in Sur are also plotted to show that they are comparable to the metabolic guilds of the S-cycle. Figure 5D shows the result of manually assigning candidate genomes in Sur to classes in terms of their ecological capabilities (see Table S7).

Out of 192 Sur genomes, 68 are known to metabolize S-compounds under culture conditions in literature reports. For instance, *Sideroxydans lithotrophicus* ES-1, a microaerophilic Fe-oxidizing bacterium has been observed to grow using thiosulfate as an energy source (Emerson *et al.*, 2013). Another 59 Sur organisms were isolated from Sulfur-rich environments such as hot springs or solfataric muds, including uncultured species with genomes assembled from metagenomic sequences. For instance, *Candidatus Desulforudis audaxviator* MP104C was isolated from basalt-hosted fluids of the deep subseafloor (Jungbluth *et al.*, 2016). Moreover, an unnamed endosymbiont of a scaly snail was sampled from a black smoker chimney (Nakagawa *et al.*, 2014). Finally, the archaon *Geoglobus ahangari* was isolated from a 2,000m depth hydrothermal vent (Manzella *et al.*, 2015). Combining these two subsets, 68% of species in Sur were confirmed by curation to be S-based.

We additionally confirmed the importance of S-cycle in gastrointestinal microbes of the genus *Campylobacter* by detecting 20 genomes with high SS values. This result is consistent with the implication of S-metabolism in the low oxygen environment of the host guts, where several inorganic (e.g., sulfates, sulfites) or organic (e.g., dietary amino acids and host mucins) S-compounds originate and are metabolized by several microorganisms. Among the microbes involved in colonic S-metabolism are SRB, many methanogens and *Campylobacter* genus (Carbonero *et al.*, 2012). Furthermore, some species of *Campylobacter* have been isolated from deep sea hydrothermal vents (Nakagawa *et al.*, 2007), suggesting that this genus plays an important role in the S cycle. The remaining species in Sur were classified in these categories: biorremediation (7), Fe-environment (2), marine (2), peatlands (2) and other environments (32).

Overall, these results highlight the broad applicability of our proposed entropy-based score to accurately predict, classify and measure the importance of the S-cycle in both in culture-derived and novel sequenced genomes without prior culture and biochemical knowledge.

### Detecting key sulfur metagenomic environments with the Sulfur Score

Encouraged by the genomic benchmark results described above, we set out to evaluate the importance of the S-cycle across 935 metagenomes in dataset Met. We calculated the *SS* for each metagenome taking into account its Mean Size Length (MSL) and the corresponding entropies calculated in dataset GenF (Table S8). The global distribution of Met is mapped in Figure 6A, with *SS* scores colored from yellow to red. To discriminate the most important S-related environments, those with *SS* values equal or larger than the 95^th^ percentile of the corresponding MSL category are shown with blue borders.

**Figure 6.**
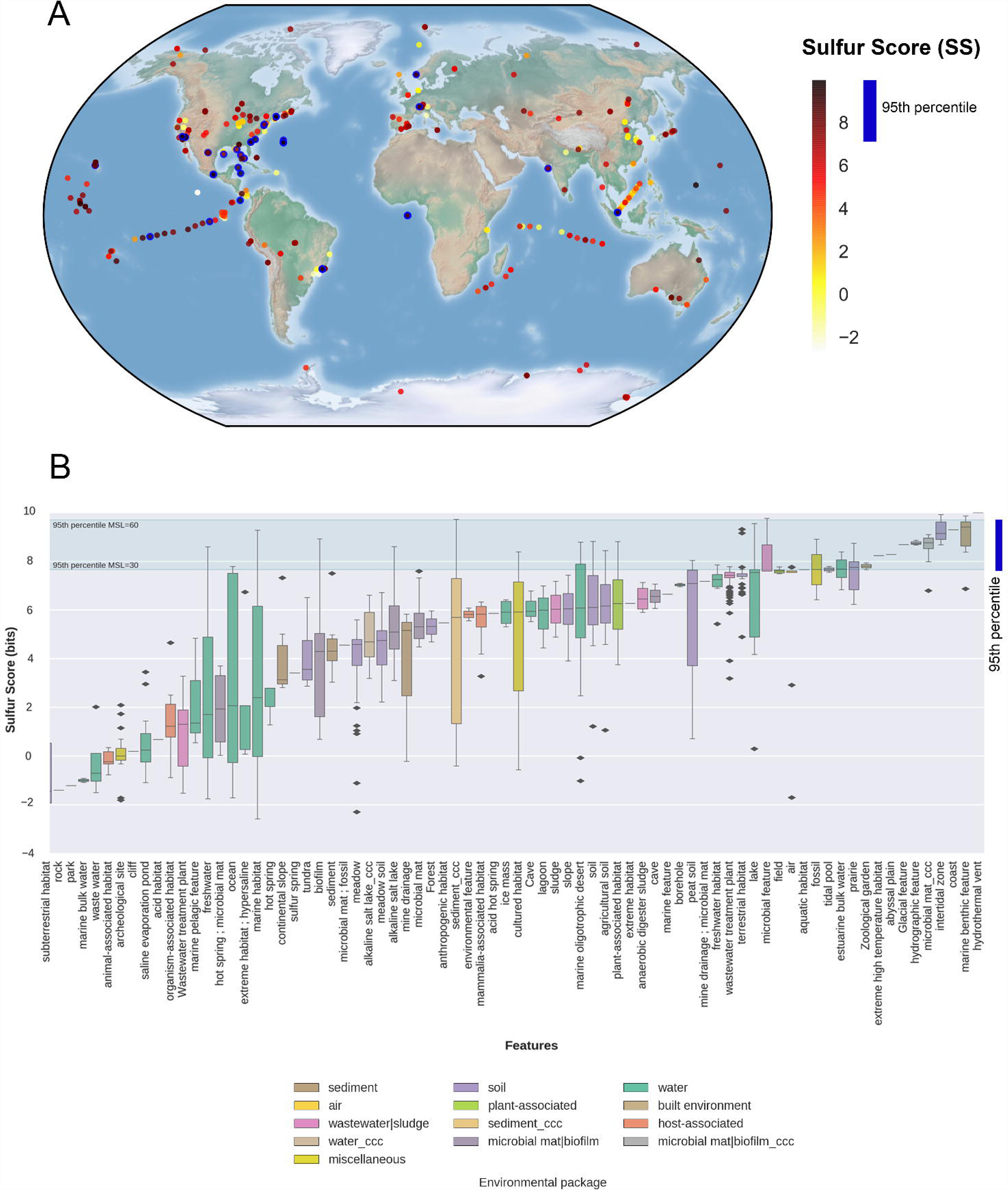
Distribution of Sulfur Score (*SS*) in the metagenomic dataset Met. A) Geo-localized metagenomes sampled around the globe are colored according to their *SS* values. The following cut-off values correspond to the 95^th^ percentiles of seven Mean Size Length classes (30, 60, 100, 150, 200, 250 and 300 aa): 7.66, 9.70, 8.81, 8.51, 8.18, 8.98 and 7.61, respectively. Circles with thick blue border indicate metagenomes with SS ≥ the 95^th^ percentile. B) Distribution of SS values observed in 935 metagenomes classified in terms of features (X-axis) and colored according to their environmental packages. Features are sorted according to their median *SS* values. ccc: metagenomes from Cuatro Cienegas, Coahuila, Mexico. Green lines indicate the lowest and largest 95^th^ percentiles observed across MSL classes.

In order to analyze some ecological patterns of the metagenomes they were further sorted by their environmental features as proposed by the Genomic Standards Consortium [GSC] and implemented in MG-RAST. Each feature corresponds to one of 13 environmental packages (EP) that standardize metadata describing particular habitats that are applicable across all GSC checklists and beyond (Field *et al.*, 2014). The EPs represent a broad and general classification containing particular features. For example, the “water” EP includes 330 metagenomes from our dataset belonging to several features such as freshwater, lakes, estuarine, marine, hydrothermal vents, etc. Each of these features has different ecological capabilities in terms of biogeochemical cycles; therefore, will likely yield different *SS* values. The results are shown in Figure 6B. In general, all the metagenomes derived from hydrothermal vents (2), marine benthic (6), intertidal (8), and our CCC microbial mats had *SS* values above the 95^th^ percentile, highlighting the importance of the S-cycle in these environments. In contrast, the metagenomes belonging to features such as sub-terrestrial habitat (7), saline evaporation pond (24) or organisms associated habitat (7) displayed consistently low or even negative *SS* values, indicating an insignificant presence of S-metabolic pathways in those environments. The remaining features have intermediate median *SS* values and contain occasionally individual metagenomes with *SS* values above the 95^th^ percentile, such as freshwater, marine, ocean or biofilm environments.

Using our approach, we identified and annotated a total of 50 high-scoring metagenomes (Table S9). According to their corresponding literature and associated metadata, all these environments can be described as Sulfur-related as they are reported to be involved in mineralization, uptake, and recycling processes of S-compounds, for example:

i. Metagenome 4525341.3 (MSL=172aa, *SS*=9.287) sequenced from costal Oligochaete worm *Olavius algarvensis*, from which metabolic pathway reconstruction revealed the presence of gamma proteobacteria symbionts that are S-oxidizing and SRB. The chemoautotrophic symbionts provide their hosts with multiple sources of nutrition such as organic carbon from autotrophic CO2 fixation driven by oxidation of reduced inorganic S-compounds (Woyke *et al.*, 2006).
ii. Metagenome 4441663.3 (MSL=158aa, *SS*=9.986) sampled from an hydrothermal vent in the Mariana Trough in 2003 (depth: 2,850 m, fluid temperature: 106°C) (Nakai *et al.*, 2011). The hydrothermal vents are highly productive ecosystems fueled by a number of reduced inorganic substances (*e.g.*, reduced S-compounds, hydrogen or methane) contained in the deep hydrothermal fluids. Through the oxidation of such compounds, chemolithoautotrophic microorganisms gain energy, which can be used for the fixation of inorganic carbon, mediating the transfer of energy from the geothermal source to higher trophic levels and thus form the basis of the unique food chains existing in these environments (Hügler *et al.*, 2010).
iii. Metagenomes 4510162.3, 4510168.3 and 4510170.3, with MSL=32, and *SS* 7.676, 7.781 and 7.772, respectively, were sampled from the marine deep-sea surface sediments around the Deep-water horizon spill in the Gulf of Mexico. They belong to ocean feature of EP sediment. The activity of key hydrocarbon degradation pathways was confirmed in these metagenomic deep-sea sediments and the presence of metabolic pathways involved in C, N and S cycles were also confirmed in the metagenomic analysis described in (Mason *et al.*, 2014).
iv. Microbial mats from CCC were also detected above the 95th percentile (MSL: 100 and *SS*: 8.945, 9.093, 9.0, and 8.978). Samples were obtained from an ultra-oligotrophic shallow lake recovered from a desiccation event due to water over-exploitation. The sequenced microbial mats showed a clear layered visual pattern structure following the S-metabolic guilds described in Figure 2A. These were assigned to EP and feature “microbial_mat_ccc”.
v. Metagenomes 4516637.3 and 4516803.3 (MSL=30, SS=7.762 and 7.753 respectively), belong to EP and feature “air”. Their high *SS* are consistent with the biogeochemical S-cycle, since the importance of gas-phase reactions of S-compounds and the formation and subsequent involvement of sulfate aerosols as cloud-forming nuclei is well established. While these reactions can be carried out without microbial intervention, it has been suggested that microbial communities might contribute to the degradation of some of these S-compounds (Cao *et al.*, 2014).

To test the reproducibility and robustness of the Sulfur Score, we conducted two further analyses. In the first one, summarized in Figure S6, we compared *SS* estimates of the Met dataset which combined Pfam entropies computed in 2014 and 2017. Despite the changes of both, the Pfam database and the Suli list, overall we find a strong correlation, yielding an *R*^2^=0.912 (Figure S6 A). A kernel density analysis of the comparison suggests a different behavior of low and high *SS* scores, with the latter being more reproducible (see Figure S6B). In the second analysis, we quantitatively tested to what extent the entropy estimates of the set of 112 Pfam domains affect the outcome of the *SS* in Gen and Met. To do so, we subsampled randomly ≈50% of those domains to compute the *SS* 1,000 times for each of the 2,017 nr-genomes and 935 metagenomes. The results, summarized in Table S10, confirm that *SS* values computed with random subsets of Pfam domains are generally lower than *SS* derived from the full list (n=112) of S-related Pfam domains in Sucy, regardless of MSL. However, as shown in Figure S7 for MSL=60, the overall ranking of metagenomes produced by computing *SS* values only with a subset of Pfam domains is broadly equivalent. Altogether, these analyses suggest that the choice of protein domains does affect the absolute scores obtained. However, high scoring metagenomes are ranked highly, even when only a reduced set of Pfam domains are employed.

The latter result confirms that a comprehensive database of protein-coding genes derived from a systematic reconstruction of global biogeochemical cycles, is necessary to apply the algorithm to quantify the importance of any given elements biogeochemical cycle in a given environment.

## CONCLUSIONS

In this study, we proposed a computational approach to address the complexity of global biogeochemical cycles on a multi-genomic scale. This methodology requires the curation of an inventory of biotic players (including genes, molecular pathways and microorganisms) and abiotic compounds involved in the cycle of interest. We focused on the S-cycle due to is importance on the biogeochemistry of the planet, but in principle this approach could be applied to any other cycle with microbial participation.

For the first time, we systematically build two databases that describe the complexity of the S-cycle based on the minimum ecosystem concept and the microbial mat model. We compiled available non-redundant fully sequenced S-based microorganisms (Suli), and all the known metabolic pathways involved in the S-cycle, taking into account also relevant abiotic S-compounds (Sucy). We used these databases as the input of a computational entropy-based approach that works in two stages. First, we used the individual entropies of protein domains annotated in Sucy to propose a list of 12 molecular markers of the S-cycle and then combined these in order to produce the Sulfur Score, a measure that informs about the presence of the molecular machinery involved in S metabolism. We benchmarked the predictive value of the Sulfur Score by producing a ROC curve, and tested its robustness with simulations against randomly subsampled subsets of the curated protein domains.

Altogether, we propose that this method can be used to evaluate other global biogeochemical cycles or complex molecular pathways across genomic and metagenomic sequence datasets, therefore allowing the detection of environmental patterns and informative samples using a single score.

## Acknowledgements

Valerie De Anda is a doctoral student from Programa de Doctorado en Ciencias Biomédicas, Universidad Nacional Autónoma de México (UNAM) and received fellowship 356832 from CONACYT. We also acknowledge funding from WWF-Alianza Carlos Slim, as well as Sep-Ciencia Básica Conacyt Proyect 238245 both to VS and LEE.

## References

Alcaraz, L.D., Olmedo, G., Bonilla, G., Cerritos, R., Hernández, G., Cruz, A., et al. (2008) The genome of Bacillus coahuilensis reveals adaptations essential for survival in the relic of an ancient marine environment. Proc. Natl. Acad. Sci. U. S. A. 105: 5803–8.

Bolhuis, H., Cretoiu, M.S., and Stal, L.J. (2014) Molecular Ecology of Microbial Mats. FEMS Microbiol. Ecol. 3: 1–16.

Canfiel, DE, Thamdrup B, Kristensen E. (2005) Aquatic Geomicrobiology, Volume 48 (Advances in Marine Biology). Elsevier Academic Press.

Cao, C., Jiang, W., Wang, B., Fang, J., Lang, J., Tian, G., et al. (2014) Inhalable Microorganisms in Beijing’s PM 2.5 and PM 10 Pollutants during a Severe Smog Event. Enviromental Sci. Technol. 48: 1499–1507.

Carbonero, F., Benefiel, A.C., Alizadeh-Ghamsari, A. H., and Gaskins, H.R. (2012) Microbial pathways in colonic sulfur metabolism and links with health and disease. Front. Physiol. 3: 1–11.

Caspi, R., Altman, T., Dreher, K., Fulcher, C. a, Subhraveti, P., Keseler, I.M., et al. (2012) The MetaCyc database of metabolic pathways and enzymes and the BioCyc collection of pathway/genome databases. Nucleic Acids Res. 40: D742–53.

Chen, L., Hu, M., Huang, L., Hua, Z., Kuang, J., Li, S., and Shu, W. (2015) Comparative metagenomic and metatranscriptomic analyses of microbial communities in acid mine drainage. ISME J. 9(7): 1579–1592.

Commenges, D. (2015) Information Theory and Statistics: an overview. ARXIV preprint arXiv:1511.00860.

Dahl (2017) Modern Topics in the Phototrophic Prokaryiotes. Metabolism, Bioenergetics and Omics. Springer International Publishing pp 27–66, 10.1007/978-3-319-51365-2_2.

Emerson, D., Field, E.K., Chertkov, O., Davenport, K.W., Goodwin, L., Munk, C., et al. (2013) Comparative genomics of freshwater Fe-oxidizing bacteria: implications for physiology, ecology, and systematics. Front. Microbiol. 4: 254.

Falkowski, P.G., Fenchel, T., and Delong, E.F. (2008) The microbial engines that drive Earth’s biogeochemical cycles. Science 320: 1034–9.

Field, D., Sterk, P., Kottmann, R., Smet, J.W. De, Amaral-zettler, L., Cole, J.R., et al. (2014) Genomic Standards Consortium Projects The Genomic Standards Consortium Initiating and Maintaining a Project within the GSC The GSC Project Description template provides a References: Stand. Genomic Sci. 599–601.

Finn, R.D., Clements, J., and Eddy, S.R. (2011) HMMER web server: interactive sequence similarity searching. Nucleic Acids Res. 39: W29–37.

Finn, R.D., Tate, J., Mistry, J., Coggill, P.C., Sammut, S.J., Hotz, H.-R., et al. (2008) The Pfam protein families database. Nucleic Acids Res. 36: D281–8.

Halevy, I., Peters, S.E., and Fischer, W.W. (2012) Sulfate burial constraints on the Phanerozoic sulfur cycle. Science 337: 331–4.

Hedges, J.I. (1992) Global biogeochemical cycles: progress and problems. Mar. Chem. 39: 67–93.

Herman EK and Kump LR (2005) Biogeochemistry of microbial mats under Precambrian environmental conditions: a modelling study. Geob 3: 77–92.

Hertz, G.Z. and Stormo, G.D. (1999) Identifying DNA and protein patterns with statistically significant alignments of multiple sequences. Bioinformatics 15: 563–77.

Hügler, M., Gärtner, A., and Imhoff, J.F. (2010) Functional genes as markers for sulfur cycling and CO2 fixation in microbial communities of hydrothermal vents of the Logatchev field. FEMS Microbiol. Ecol. 73: 526–537.

Jones, P., Binns, D., Chang, H.Y., Fraser, M., Li, W., McAnulla, C., et al. (2014) InterProScan 5: Genome-scale protein function classification. Bioinformatics 30: 1236–1240.

Jungbluth SP, Glavina del Rio T, Tringe SG, Stepanauskas R, Rappé MS. (2016) Genomic comparisons of a bacterial lineage that inhabits both marine and terrestrial deep subsurface systems. PeerJ Preprints 4:e2592v1https://doi.org/10.7287/peerj.preprints.2592v1

Kanehisa, M. and Goto, S. (2000) KEGG: kyoto encyclopedia of genes and genomes. Nucleic Acids Res. 28: 27–30.

Khodadad, C. L.M. and Foster, J.S. (2012) Metagenomic and metabolic profiling of nonlithifying and lithifying stromatolitic mats of Highborne Cay, The Bahamas. PLoS One 7: e38229.

Kullback, S. and Leibler, R.A. (1951) On Information and Sufficiency. Ann. Math. Stat. 22: 79–86.

Li, Y., Yu, S., Strong, J., and Wang, H. (2012) Are the biogeochemical cycles of carbon, nitrogen, sulfur, and phosphorus driven by the “FeIII–FeII redox wheel” in dynamic redox environments? J. Soils Sediments 12: 683–693.

Llorens-Marès, T., Yooseph, S., Goll, J., Hoffman, J., Vila-Costa, M., Borrego, C. M., et al. (2015) Connecting biodiversity and potential functional role in modern euxinic environments by microbial metagenomics. ISME J. 9(7): 1579–92.

Loy, A., Ku, K., Lehner, A., Drake, H.L., and Wagner, M. (2004) Microarray and Functional Gene Analyses of Sulfate-Reducing Prokaryotes in Low-Sulfate, Acidic Fens Reveal Cooccurrence of Recognized Genera and Novel Lineages. Appl. Environ. Microbiol. 70: 6998–7009.

Magrane, M. and Consortium, U.P. (2011) UniProt Knowledgebase: A hub of integrated protein data. Database 2011: 1–13.

Manzella, M.P., Holmes, D.E., Rocheleau, J.M., Chung, A., Reguera, G., and Kashefi, K. (2015) The complete genome sequence and emendation of the hyperthermophilic, obligate iron-reducing archaeon “Geoglobus ahangari” strain 234T. Stand. Genomic Sci. 10: 77.

Mason, O.U., Scott, N.M., Gonzalez, A., Robbins-Pianka, A., Bælum, J., Kimbrel, J., et al. (2014) Metagenomics reveals sediment microbial community response to Deepwater Horizon oil spill. ISME J. 8: 1464–75.

Meyer, B. and Kuever, J. (2007) Molecular analysis of the diversity of sulfate-reducing and sulfur-oxidizing prokaryotes in the environment, using aprA as functional marker gene. Appl. Environ. Microbiol. 73: 7664–79.

Morales, S. E. and Holben, W.E. (2011) Linking bacterial identities and ecosystem processes: Can “omic” analyses be more than the sum of their parts? FEMS Microbiol. Ecol. 75: 2–16.

Moreno-Hagelsieb, G., Wang, Z., Walsh, S., and ElSherbiny, A. (2013) Phylogenomic clustering for selecting non-redundant genomes for comparative genomics. Bioinformatics 29: 947–9.

Nakagawa, S., Shimamura, S., Takaki, Y., Suzuki, Y., Murakami, S., Watanabe, T., et al. (2014) Allying with armored snails: the complete genome of gammaproteobacterial endosymbiont. ISME J. 8: 40–51.

Nakagawa, S., Takaki, Y., Shimamura, S., Reysenbach, A.-L., Takai, K., and Horikoshi, K. (2007) Deep-sea vent epsilon-proteobacterial genomes provide insights into emergence of pathogens. Proc. Natl. Acad. Sci. U. S. A. 104: 12146–12150.

Nakai, R., Abe, T., Takeyama, H., and Naganuma, T. (2011) Metagenomic Analysis of 0.2-μm-Passable Microorganisms in Deep-Sea Hydrothermal Fluid. Mar. Biotechnol. 13: 900–908.

Newman, D.K. and Banfield, J.F. (2002) Geomicrobiology: how molecular-scale interactions underpin biogeochemical systems. Science 296: 1071–7.

Odum EP (1993) Ecology and our endangered life-support systems, 2nd edn. Sinauer Associates Inc., Sunderland, Massachusetts. 301 pp.

Olson, K.R., Straub, K.D., and Straub, K.D. (2016) The Role of Hydrogen Sulfide in Evolution and the Evolution of Hydrogen Sulfide in Metabolism and Signaling The Role of Hydrogen Sulfide in Evolution and the Evolution of Hydrogen Sulfide in Metabolism and Signaling. Physiology 31: 60–72.

Pedroni, P., Volpe, A.D., Galli, G., Mura, G.M., Pratesi, C., and Grandi, G. (1995) Characterization of the locus encoding the [Ni-Fe] sulfhydrogenase from the archaeon Pyrococcus furiosus: Evidence for a relationship to bacterial sulfite reductases. Microbiology 141: 449–458.

Pinckney, J. L. and Paerl, H.W. (1997) Anoxygenic photosynthesis and nitrogen fixation by a microbial mat community in a bahamian hypersaline lagoon. Appl. Environ. Microbiol. 63: 420–6.

Quaiser, A., Zivanovic, Y., Moreira, D., and López-García, P. (2011) Comparative metagenomics of bathypelagic plankton and bottom sediment from the Sea of Marmara. ISME J. 5: 285–304.

Rabus, R., Hansen, T., and Widdel, F. (2013) “Dissimilatory sulfate- and sulfur-reducing prokaryotes,” in The Prokaryotes, eds E. Rosenberg, E. Delong, S. Lory, E. Stackebrandt, and F. Thompson. Heidelberg: Springer. 309–404.

Santos, A.A., Venceslau, S.S., Grein, F., Leavitt, W.D., Dahl, C., Johnston, D.T., and Pereira, I.A.C. (2015) A protein trisulfide couples dissimilatory sulfate reduction to energy conservation. Science (80-.). 350: 1541–1545.

Stewart, F.J., Dmytrenko, O., Delong, E.F., and Cavanaugh, C.M. (2011) Metatranscriptomic analysis of sulfur oxidation genes in the endosymbiont of solemya velum. Front. Microbiol. 2: 134.

Swingley, W.D., Meyer-Dombard, D.R., Shock, E.L., Alsop, E.B., Falenski, H.D., Havig, J.R., and Raymond, J. (2012) Coordinating environmental genomics and geochemistry reveals metabolic transitions in a hot spring ecosystem. PLoS One 7: e38108.

Taguchi , Y., Sugishima, M., and Fukuyama, K. (2004) Crystal Structure of a Novel Zinc-Binding ATP Sulfurylase from Thermus. Biochemistry 43: 4111–4118.

Tu, Q., Yu, H., He, Z., Deng, Y., Wu, L., Van Nostrand, J.D., et al. (2014) GeoChip 4: A functional gene-array-based high-throughput environmental technology for microbial community analysis. Mol. Ecol. Resour. 14: 914–928.

Widder, S., Allen, R.J., Pfeiffer, T., Curtis, T.P., Wiuf, C., Sloan, W.T., et al. (2016) Challenges in microbial ecology: building predictive understanding of community function and dynamics. ISME J. 10: 2557–2568.

Woyke, T., Teeling, H., Ivanova, N.N., Huntemann, M., Richter, M., Gloeckner, F.O., et al. (2006) Symbiosis insights through metagenomic analysis of a microbial consortium. Nature 443: 950–5.

Zhao, M., Xue, K., Wang, F., Liu, S., Bai, S., Sun, B., et al. (2014) Microbial mediation of biogeochemical cycles revealed by simulation of global changes with soil transplant and cropping. ISME J. 8(10): 2045–55.

